## Supplementary Table for "Coupled protein synthesis and ribosome-guided piRNA processing on mRNAs"

**Supplementary Materials**

Materials and Methods

Table S1

Figs. S1 to S7

**Extended Data information for:**

### Coupled protein synthesis and ribosome-guided piRNA processing on mRNAs

Yu H. Sun, Ruoqiao Huiyi Wang, Khai Du, Jihong Zheng, Li Huitong Xie, Amanda A. Pereira, Chao Zhang, Emiliano P. Ricci, and Xin Zhiguo Li

**This PDF file includes:**

Caption for Additional Data Table S1

Figs. S1 to S7

**Other Supplemental Materials for this manuscript includes the following:**

Table S1 (separate file)

Detailed information and statistics for the sequencing data used in this study.

**Additional Data Table S1 (separate file)**

Detailed information and statistics for the sequencing data used in this study (species refers to numbers of unique sequences in a set of reads).

(A) Ribo-seq statistics: reads and species

(B) Small RNA sequencing statistics: reads and species.

(C) RNA-seq statistics: reads and species.

(C) Degradome-seq statistics: reads and species.

(E) Genome coordinates for the 30 3´UTR piRNA precursors with annotated ORFs provided in UCSC BED format (i.e., 0-based) for mm10.

(F) Genome coordinates for the 43 control mRNAs with annotated ORFs provided in UCSC BED format (i.e., 0-based) for mm10.

(G) Genome coordinates for the 23 piRNA precursor mRNAs with annotated ORFs provided in UCSC BED format (i.e., 0-based) for galgal6.

(H) Genome coordinates for the 23 uppl chicken homolog mRNAs with annotated ORFs provided in UCSC BED format (i.e., 0-based) for galgal6.

#### EXTENDED DATA FIGURE LEGENDS

**Fig. S1, Related to Fig. 1. A-MYB regulates 3´UTR piRNA precursor transcription and MOV10L1 regulates their processing.** (A) Top, boxplots represent 3´UTR piRNA density per gene as spermatogenesis progresses in reads per kilobase per million mapped reads (rpkm). Bottom, boxplots represent uppl transcript expression, as measured by RNA-seq in transcripts per million (tpm), at each of the five wild-type development stages examined. (B) Transcript and piRNA abundance in heterozygous (Het) and homozygous *A-Myb* (Mut) point mutant testes are shown for an illustrative example of uppl mRNA at 14.5 and 17.5 dpp: the 3´UTR piRNA gene *pi-Mlc1.1*. Ppm, parts per million reads mapped to the genome. (C) Histology of adult testis sections from *Mov10l1*$^{CKO/\Delta}$ *Neurog3-cre* (*top*) and *Mov10l1*$^{CKO/\Delta}$ (*bottom*). RS, round spermatids; ES, elongating spermatids. (D) Aggregated data for RNA abundance from *Mov10l1*$^{CKO/\Delta}$ testes (*left*) and *Mov10l1*$^{CKO/\Delta}$ *Neurog3-cre* (*right*) across 5´UTRs, ORFs, and 3´-UTRs of the control mRNAs from adult testes. The x-axis shows the median length of these regions, and the y-axis represents the 10% trimmed mean of relative abundance. Ppm, parts per million. (E) Boxplots of RNA abundance ratios of *Mov10l1*$^{CKO/\Delta}$ *Neurog3-cre* versus *Mov10l1*$^{CKO/\Delta}$ adult testis (here and elsewhere, uppl mRNAs in green and control mRNAs in black). (F) Boxplots of intron RNA abundance ratios of *Mov10l1*$^{CKO/\Delta}$ *Neurog3-cre* versus *Mov10l1*$^{CKO/\Delta}$ adult testis. (G) Aggregated data for MOV10L1 CLIP-seq abundance (10% trimmed mean) on uppl mRNAs (left) and control mRNAs (right) from wild-type testis[13].

**Fig. S2, Related to Fig. 2. uppl mRNAs are bifunctional, coding for proteins and piRNAs.** (A) Boxplots showing the density of uniquely mapping piRNAs in the exons, introns, and 100 nt after the 3´end (i.e., the polyadenylation site, PAS) of uppl mRNAs. (B) Boxplots showing unspliced transcript length distributions. (C) Boxplots showing number of introns. (D) Boxplots showing exon length distributions. (E) Boxplots showing codon adaptation index (CAI)[69,70]. (F) Boxplots showing normalized codon adaptation index (CAI)[152].

**Fig. S3, Related to Fig. 3. Ribosomes serve as templates for endonucleolytic cleavage to form 3´UTR piRNA 5´ends.** (A) Metagene plots of RPFs at 5´UTR, ORF, and 3´UTR of the control mRNAs in adult testes. The x-axis shows the median length of these regions, and the y-axis represents the mean of normalized abundance. Ppm, parts per million. (B) Boxplots of the RPF abundance per mRNA at ORF and 3´UTR in adult testes. (C) Boxplots of 3´UTR piRNA abundance per uppl mRNA with and without RNase A & T1 treatment in adult wild-type testes. p value was determined by paired Wilcoxon rank sum test. (D) Boxplots of RFP abundance in anti-HA IP versus input (before IP) ($n$ = 3). The Ribo-seq libraries before (input) and after anti-HA IP were normalized to reads mapping to mRNA ORFs in adult RiboTag testes. (E) Boxplots of distance spectra of 5´-ends of RPFs from adult testis that overlap simulated sequences. The 5´-end overlap analyses between RPFs and simulated sequences were computed 10,000 times. (F) Boxplots of distance spectra of 5´-ends of sheared RNA fragments that overlap piRNAs of adult testes.

**Fig. S4, Related to Fig. 4. Biphasic piRNA biogenesis at ORFs and 3´UTRs.** (A) Schematic of the modified ribosome profiling library construction procedure which captures 5´OH RPF (*left*) and 5´P RPFs (*right*). (B) Sequence logos depicting nucleotide bias at 5´-ends and 1 nt upstream of 5´-ends. Top to bottom: anti-HA immunoprecipitated RPF species with a 5´monophosphate end, and anti-HA immunoprecipitated RPF species with a 5´OH end, from adult testes. (C) Boxplots of the length of all putative ORFs embedded in each uppl 3´ UTR. (D) Boxplots of change in RPF abundance with harringtonine treatment ($n$ = 2) versus no treatment adult wild-type testes. (E) Boxplots of the ratios of RNA abundance (normalized with length) at 3´UTRs versus RNA abundance at ORFs of uppl mRNAs in adult wild-type testes. (F) Distance spectrum of 5´-ends of anti-HA immunoprecipitated 5´P RNA species from ORFs (*left*) and from 3´UTRs (*right*) that overlap piRNAs in adult testes ($n$ = 3). Data are mean ± standard deviation. (G) Metagene analysis of piRNA abundance from 400 nt upstream stop codon to 1,000 nt downstream stop codon from adult wild-type testes. Ppm, parts per million. (H) A general model of ribosome-guided piRNA biogenesis. The grey and tan bubbles represent the large and small subunits of ribosomes respectively. The blue bubbles represent PIWI proteins. Grey circle in PIWI proteins on the left represents MID domain that recognizes the 5′-phosphate (5′P), on the right represents PAZ domain, in between represents PIWI domain. piRNA precursors are synthesized by RNA polymerase II, and contain the 5′-cap, exons, introns, and a poly(A) tail. The transcription of pachytene piRNA genes and 3′UTR piRNA gene is controlled by A-MYB. The PLD6-mediated cleavage before uridine generates strings of head-to-tail phased piRNAs that will be further trimmed and methylated. Ribosomes translate the piRNA precursors in a canonical fashion through initiating at the start codon near the 5′-cap. Post-termination ribosomes traverse in the 3′ direction. The 5′-end-loaded PIWI protein and the downstream ribosome specify the endonuclease PLD6 cleavage site, and determine the 5′- and 3′-ends of head-to-tail strings before 3′-end trimming. piRNA biogenesis at the 3′ UTRs requires TDRD5 protein. Transcripts containing ORFs are processed to piRNAs inefficiently that does not require TDRD5 (Gold arrows). This model applies to precursor source of both lncRNAs and mRNAs.

**Fig. S5, Related to Fig. 5. Co-translational piRNA processing.** (A) miRNA abundance in *A-Myb* mutant and heterozygous testes at 14.5 dpp. (sample size $n = 2$ independent biological samples). Data are mean ± standard deviation. (B) A-MYB ChIP-seq and input signal near the two miRNA genes that are decreased in *A-Myb* mutant. For each gene, the figure also reports the miRNA abundance in *A-Myb* mutant and heterozygous testes at 17.5 dpp. (C) Boxplots of the changes of RPF abundance in *Mov10l1* CKO mutants ($n = 3$) compared to littermate controls in adult testes.

**Fig. S6, Related to Fig. 6. Conserved piRNA biogenesis from mRNA 3′UTRs.** (A) The distance from the annotated transcription start site of each uppl mRNA gene (middle) and from the 5′-ends of uppl mRNA 3′UTRs (right) to the nearest H3K4me3 peak, and from the annotated transcription start site of each uppl mRNA gene to the nearest A-MYB peak (left) in roosters. (B) Boxplots of piRNA abundance per mRNA in adult wild-type mouse testes. (C) Boxplots of piRNA abundance per mRNA in adult rooster testes. (D) Residual plot of partial correlation between piRNA and RPF abundance conditional on RNA-seq abundance at the chicken uppl

3´UTR (left) and uppl ORF (right) in adult rooster testes. Linear regression was performed between piRNA and RNA-seq abundance, RPF and RNA-seq abundance on uppl mRNA 3´UTRs separately, and residuals were plotted with smooth parameter 'method=lm'. (E) Boxplots of distance spectra of 5´-ends of RPFs from adult rooster testis that overlap simulated sequences. The 5´-end overlap analyses between RPFs and simulated sequences were computed 10,000 times.

**Extended Data Fig. 7, Related to Fig. 7. piRNA precursor mRNAs contain transposon fragments.** (A) The distance from the uppl mRNA gene and control mRNA gene to the nearest annotated TE superfamilies in mouse genome. (B) Boxplots showing the fraction of transcript intron sequence correspondence to sense (blue) and anti-sense (red) transposon sequences in mouse genome. (C) Boxplots showing the fraction of transcript exon sequence correspondence to sense (blue) and antisense (red) SINE sequences in mouse genome. (D) Sequence logo showing the nucleotide composition of sense SINE-piRNA species that uniquely map to mouse uppl mRNAs. (E) The 5′-5′ overlap between piRNAs from opposite strands was analyzed to determine if sense SINE piRNAs from mouse uppl mRNAs display Ping-Pong amplification in trans. The number of pairs of piRNA reads at each position is reported. The *Z*-score indicates that a significant ten-nucleotide overlap ("Ping-Pong") was detected. *Z*-score >1.96 corresponds to $p$-value < 0.05. (F) Boxplots showing the fraction of transcript exon sequence correspondence to sense (blue) and antisense (red) LINE sequences in chicken genome.
